# Human paternal and maternal demographic histories: insights from high-resolution Y chromosome and mtDNA sequences

**DOI:** 10.1101/001792

**Authors:** Sebastian Lippold, Hongyang Xu, Albert Ko, Mingkun Li, Gabriel Renaud, Anne Butthof, Roland Schröder, Mark Stoneking

## Abstract

To investigate in detail the paternal and maternal demographic histories of humans, we obtained ∼500 kb of non-recombining Y chromosome (NRY) sequences and complete mtDNA genome sequences from 623 males from 51 populations in the CEPH Human Genome Diversity Panel (HGDP). Our results: confirm the controversial assertion that genetic differences between human populations on a global scale are bigger for the NRY than for mtDNA; suggest very small ancestral effective population sizes (<100) for the out-of-Africa migration as well as for many human populations; and indicate that the ratio of female effective population size to male effective population size (N_f_/N_m_) has been greater than one throughout the history of modern humans, and has recently increased due to faster growth in N_f_. However, we also find substantial differences in patterns of mtDNA vs. NRY variation in different regional groups; thus, global patterns of variation are not necessarily representative of specific geographic regions.

---

Comparisons of mtDNA and NRY variation have provided numerous important insights into the maternal and paternal histories of human populations^1–3^. However, such comparisons are limited by methodological differences in how mtDNA and NRY variation have been typically assayed. MtDNA variation is usually investigated by sequencing hypervariable segments of the control region, (or, increasingly, via complete mtDNA genome sequences), while human NRY variation is routinely assayed by genotyping SNPs of interest, often in combination with Y-STR loci. Nevertheless, NRY SNP-typing has several drawbacks due to the ascertainment bias inherent in the selection of SNPs^1, 4, 5^. This ascertainment bias precludes many analyses of interest, such as dating the age of the NRY ancestor or particular divergence events in the NRY phylogeny, as well as demographic inferences such as population size changes. Moreover, the difference in molecular methods used to assay NRY vs. mtDNA variation can complicate the interpretation of differences between patterns of NRY and mtDNA variation. For example, the seminal finding that NRY differences are bigger than mtDNA differences among global populations of humans, and that this is due to a higher rate of female than male migration due to patrilocality^6^, may instead reflect methodological differences in how mtDNA vs. NRY variation is assayed^7^. Another fundamental question concerns whether or not male and female effective population sizes have been the same over time. Attempts to address this question using the ratio of X chromosome to autosomal DNA diversity have come up with conflicting answers^8, 9^, which may in part reflect the use of different methods that capture information about effective population size at different times in the past^10^. Moreover, the ratio of X to autosome diversity varies along the X chromosome, depending how far polymorphic sites are from genes ^11^, indicating a potential role for selection in distorting effective population size estimates from comparisons of X chromosome to autosomal DNA diversity. These questions – namely, are genetic differences between populations and effective population sizes the same for males and females –as well as other fundamental aspects of human maternal and paternal demographic history remain unanswered.

Recently, analyses have been carried out of NRY sequences obtained as part of whole genome sequencing projects^12–14^. While these studies provide very detailed insights into NRY diversity, they are nonetheless limited by the expense of whole genome sequencing, which precludes comprehensive global sampling. To allow for more accurate comparisons between mtDNA and NRY variation and to permit demographic inferences based on the NRY, we developed a capture-based array to enrich Illumina sequencing libraries for ∼500 kb of NRY sequence (**Supplementary Table 1**). We used this approach to obtain NRY sequences from 623 males in the HGDP^15^, from 51 globally-distributed populations. We also obtained complete mtDNA genome sequences from the entire HGDP, allowing us to investigate and directly compare the paternal and maternal relationships of global human populations in unprecedented detail.

The average coverage of the NRY sequences was 14.5X (range 5X-37.5X, **Supplementary Figure 1**), while for the mtDNA genome sequences the average coverage was 640X (range 46X-4123X, **Supplementary Figure 1**). After quality-filtering, imputation, and removal of sites with a high number of recurrent mutations, there remained 2228 SNPs in the NRY sequences. The mtDNA analyses here are restricted to the 623 males for which NRY sequences were obtained, for which tere were 2163 SNPs; results based on the mtDNA genome sequences from the entire HGDP (952 individuals) did not differ from those based on the subset of 623 males (**Supplementary Figure 2**). More details about the results from each individual, including mtDNA and NRY haplogroups, are provided in **Supplementary Table 2**. The mtDNA sequences have been deposited in Genbank with accession numbers KF450814 -KF451871. The NRY raw data are in the European Nucleotide Archive (ENA) (http://www.ebi.ac.uk/ena/home) with the study accession number PRJEB4417 (sample accession numbers ERS333252-ERS333873).

Basic summary statistics for the mtDNA and NRY diversity in each population are provided in **Supplementary Table 3**. As the sample sizes for many of the individual populations are quite small, for most subsequent analyses we grouped the populations into the following regions (based on analyses of genome-wide SNP data^16, 17^): Africa, America, Central Asia, East Asia, Europe, Middle East/North Africa (ME/NA), and Oceania (the regional affiliation for each population is in **Supplementary Table 2**). The Adygei, Hazara, and Uygur were excluded from these groupings due to extensive admixture evident in the genome-wide SNP data^16, 17^. We stress that the use of regional names is a convenience to refer to these groupings of these specific populations, and should not be taken to represent the entirety of the regions (e.g., “Africa” refers to the results based on the analysis of the combined African HGDP samples, not to Africa in general).

Some basic summary statistics concerning mtDNA and NRY diversity for the regions are provided in **Table 1**. Notably, there is substantial variation among regions in amounts of mtDNA vs. NRY diversity; this is shown further in the comparison of the mean number of pairwise differences (mpd) for mtDNA and the NRY (**Figure 1a**). The mtDNA mpd for Africa is about twice that for other regions, in keeping with previous observations of substantially greater mtDNA diversity in Africa than outside Africa^18, 19^. However, the NRY mpd is greatest in the Middle East/North Africa region, and only slightly greater in Africa than in the other regions (with the exception of the Americas, which show substantially lower NRY diversity). The greater NRY diversity in the Middle East/North Africa region can be attributed to male-biased admixture involving substantially-diverged NRY haplogroups. Nevertheless, there are striking differences in the ratio of NRY:mtDNA mdp (**Table 1**), with Africa, Central Asia, and the Americas having significantly less NRY diversity relative to mtDNA diversity, compared to the other regional groups. These results indicate substantial regional variation in the maternal vs. paternal demographic history of human populations.

**Table 1.** Summary statistics for regional groups. n, sample size; H, number of different haplotypes (sequences); S, number of polymorphic sites; mpd ± SE, mean number of pairwise differences ± standard error; π ± SE, nucleotide diversity ± standrad error; mpd ratio, ratio of the mpd_NRY_/mpd_mtDNA_. * group ratios that differ significantly (p < 0.05) from the overall average ratio for the entire HGDP, based on random resampling of NRY and mtDNA sequences.

| -----NRY----- |  |  |  |  |  | -----mtDNA----- |  |  |  |  |
| --- | --- | --- | --- | --- | --- | --- | --- | --- | --- | --- |
| Group | n | H | S | mpd ± SE | $\pi$ ± SE<sup>a</sup> | H | S | mpd ± SE | $\pi$ ± SE<sup>b</sup> | mpd ratio |
| Africa | 85 | 71 | 545 | 41.0 ± 18.0 | 80 ± 40 | 70 | 617 | 78.3 ± 34.0 | 47 ± 23 | 0.52* |
| CentralAsia | 146 | 106 | 524 | 32.1 ± 14.1 | 62 ± 31 | 131 | 833 | 42.4 ± 18.5 | 26 ± 12 | 0.76* |
| EastAsia | 162 | 141 | 709 | 35.0 ± 15.3 | 71 ± 36 | 156 | 899 | 42.3 ± 18.5 | 26 ± 12 | 0.83 |
| ME/NA | 75 | 47 | 301 | 42.7 ± 18.7 | 85 ± 40 | 71 | 618 | 42.0 ± 18.4 | 25 ± 12 | 1.02 |
| Europe | 79 | 68 | 350 | 30.0 ± 13.2 | 58 ± 31 | 78 | 432 | 29.3 ± 12.9 | 18 ± 9 | 1.02 |
| Oceania | 17 | 16 | 147 | 34.7 ± 15.9 | 71 ± 36 | 16 | 175 | 41.9 ± 19.2 | 25 ± 13 | 0.83 |
| America | 22 | 19 | 96 | 11.8 ± 5.5 | 22 ± 13 | 15 | 148 | 34.9 ± 15.8 | 21 ± 11 | 0.39* |

<sup>a</sup> multiply values by 10<sup>-6</sup>
<sup>b</sup> multiply values by 10<sup>-4</sup>

**Figure 1.**
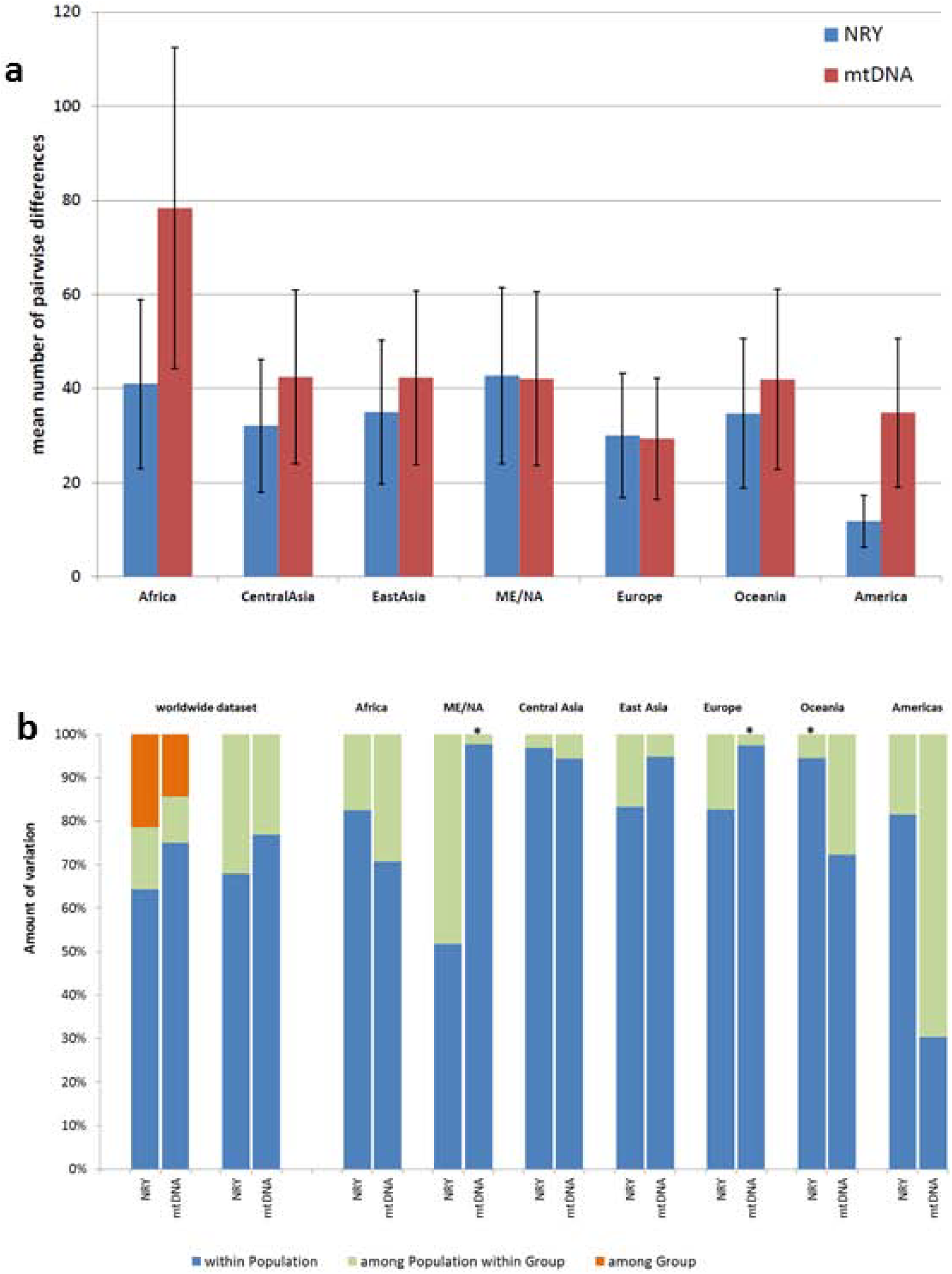
Diversity and AMOVA results. (**a**) Mean number of pairwise differences (and SE bars) for the NRY and mtDNA sequences from each regional group. (**b**) AMOVA results for the entire worldwide dataset, and for each regional group of populations. Two comparisons are shown for the entire dataset; the left comparison includes regional groups as an additional hierarchical level, while the right one does not. * indicates that the among population component of diversity does not differ significantly from zero (after Bonferroni adjustment of the p-value for multiple comparisons).

An outstanding question is whether or not there are differences in the relative amounts of between-population vs. within-population diversity for mtDNA vs. the NRY. Some studies have found much larger between-population differences for the NRY than for mtDNA^6^, while others have not^7^. To address this question, we carried out an AMOVA; the results (**Figure 1b**) show that in the entire worldwide dataset, the between-population differences are indeed bigger for the NRY (∼36% of the variance) than for mtDNA (∼25% of the variance). However, there are substantial differences among the regional groups. The ME/NA, East Asia, and Europe regional groups follow the worldwide pattern in having bigger between-population differences for the NRY than for mtDNA. In contrast, Africa, Oceania, and the Americas have substantially bigger between-population differences for mtDNA than for the NRY, while for central Asia the between-population variation is virtually identical for the NRY and mtDNA. These regional differences likely reflect the influence of sex-biased migrations and admixture, and moreover indicate that focussing exclusively on the worldwide pattern of mtDNA vs. NRY variation misses these important regional differences.

Multidimension-scaling (MDS) plots based on Φ_ST_ distances further indicate regional differences in mtDNA vs. NRY variation (**Figure 2**). Nonetheless, and despite the small sample sizes at the population level, both mtDNA and NRY Φ_ST_ distances are significantly correlated with geographic distances between populations (Mantel tests with 1000 replications: mtDNA, r = 0.41, p < 0.001; NRY, r = 0.36, p = 0.002) as well as with each other (r = 0.23, p = 0.025). Thus, NRY and mtDNA diversity are both highly associated with geographic distances among populations.

**Figure 2.**
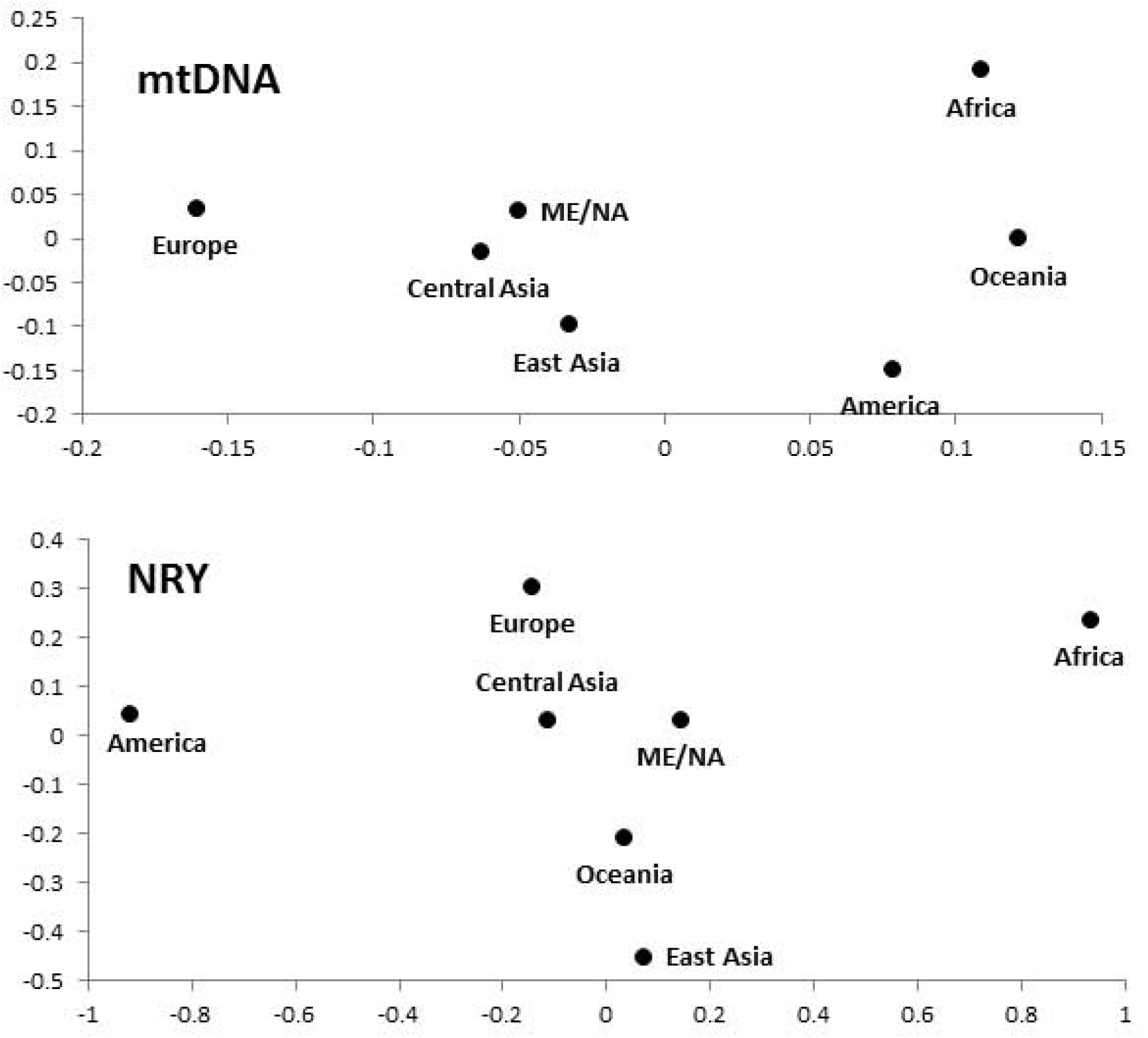
MDS plots based on Φ_ST_ distances among regional groups. The stress values are 0.055 for the mtDNA plot and 0.031 for the NRY plot.

We used a Bayesian method to estimate the phylogeny and divergence times for both mtDNA and the NRY (**Supplementary Figure 2**); for the latter, we used both a “fast” mutation rate of 1 × 10^−9^/bp/year and a “slow” mutation rate of 0.62 × 10^−9^/bp/year as there is currently much uncertainty regarding mutation rates^20–22^. The resulting phylogenies are quite consistent with the existing mtDNA and NRY phylogenies^23, 24^, with some small discrepancies involving lineages that are not well-resolved. The age of the mtDNA ancestor is estimated to be about 160 thousand years (ky), and the ages of the non-African mtDNA lineages M and N are about 65-70 ky, in good agreement with previous estimates^18^. Our estimate for the age of the NRYancestor is 103 ky based on the fast rate, and 165 ky based on the slow rate; however these estimates do not include the recently-discovered “A00” lineage^20^, which would result in much older ages for the NRY ancestor. The close agreement between the slow NRY ancestor age (165 ky) and the mtDNA ancestor age (160 ky) might be taken as evidence in favor of the slow NRY mutation rate. However, the slow NRY mutation rate gives an estimated age for the intial out-of-Africa divergence of about 100 ky, and an age for the divergence of Amerindian-specific haplogroup Q lineages of about 20 ky, while the fast rate gives corresponding estimates of about 60 ky for out-of-Africa and about 12.5 ky for Amerindian haplogroup Q lineages, in better agreement with the mtDNA and other evidence for these events^18, 25–27^. Given the current uncertainty over mutation rate estimates, we have chosen to use either both estimates in further analyses (e.g., Bayesian skyline plots) or an average of the fast and slow rates (e.g., in simulation-based analyses).

NRY and mtDNA haplogroup frequencies per population are shown in **Supplementary Table 4**. NRY haplogroups for the CEPH-HGDP males were previously determined by SNP-genotyping^28^, and the phylogenetic relationships for the NRY sequences are generally concordant with the SNP-genotyping results (with some exceptions, discussed in the legends to **Supplementary Figures 4-13**). The haplogroup frequencies provide further insights into some of the different regional patterns of mtDNA vs. NRY diversity noted previously, and these are discussed in the legend to **Supplementary Table 4**. We note some additional features in the individual NRY haplogroup phylogenies provided in **Supplementary Figures 4-13**, while the full mtDNA phylogeny is provided in **Supplementary Figure 14**.

Sequence-based analysis of NRY variation permits demographic analyses that cannot be carried out with ascertained SNP genotype data. As an example, we estimated the history of population size changes via Bayesian skyline plots (BSPs) for the NRY and mtDNA genome sequences for each region (**Figure 3**). These results should be interpreted cautiously, both because of the small sample sizes for some of the regions (in particular, America and Oceania), and because grouping populations with different histories can produce spurious signals of population growth^29^. Nevertheless, both the mtDNA and NRY BSPs indicate overall population growth in almost all groups, but for mtDNA there is a more pronounced signal of growth at around 15,000-20,000 years ago than there is for the NRY, and during much of the past it appears as if the effective size for females was larger than that for males.

**Figure 3.**
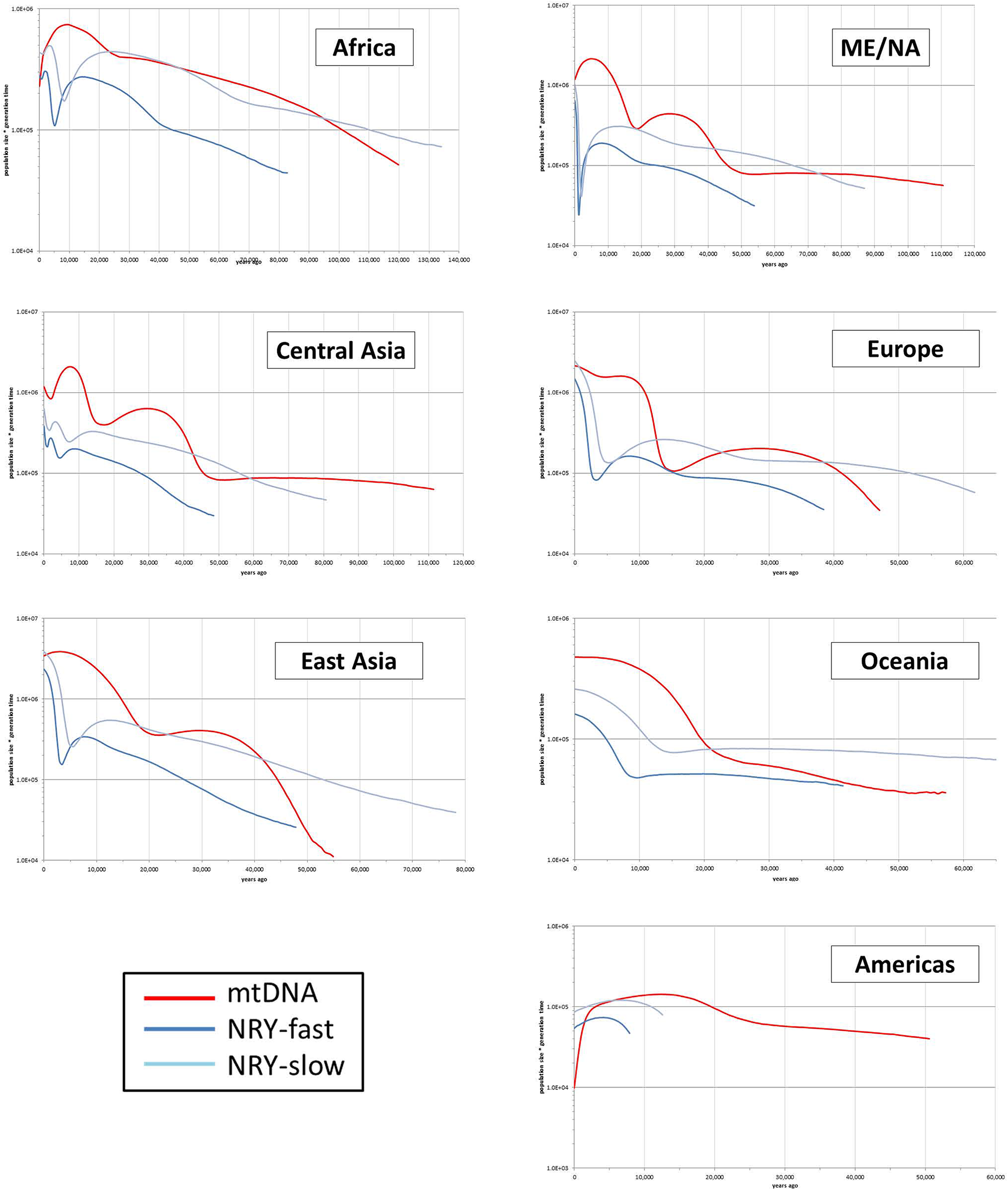
Bayesian skyline plots for regional groups. Two curves are shown for the NRY data, based on “fast” and “slow” mutation rate estimates.

To further investigate female and male demographic history, we used a simulation-based approach (**Supplementary Figure 15**) to estimate the current and ancestral effective population size for females (N_f_) and males (N_m_) for Africa, Europe, East Asia, Central Asia, Oceania, and the Americas (excluding ME/NA populations because of their admixed history). We also estimated the ancestral N_f_ and N_m_ for the out-of-Africa migration. The detailed results are provided in **Supplementary Figures 16-19** and **Supplementary Tables 5-10**; a pictorial depiction of the results is in **Figure 4**. These results indicate a small founding size in Africa of about 60 females and 30 males (all population sizes are effective population sizes); migration out of Africa about 75 ky ago (kya) associated with a bottleneck of around 25 females and 15 males; migrations from this non-African founding population to Oceania 61 kya, to Europe 49 kya, to Central and East Asia 37 kya, and from East Asia to the Americas about 15 kya. There was concomitant population growth in all regions (with the most growth in East Asia); however, throughout history the mtDNA and NRY results indicate consistently larger effective population sizes for females than for males (except, possibly, in the ancestors of East Asians).

**Figure 4.**
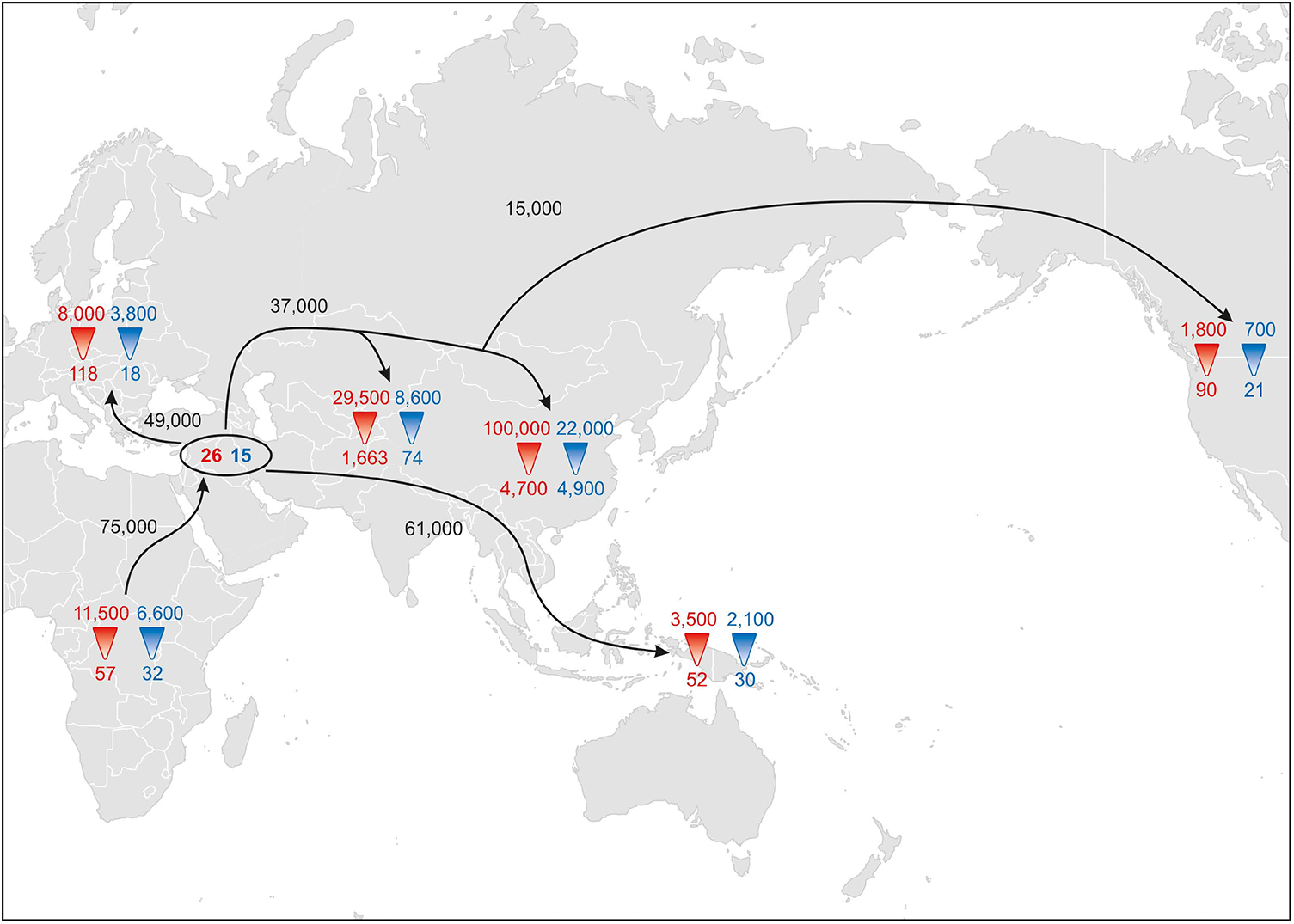
Pictorial representation of the simulation results estimating divergence times and female and male effective population sizes. Red numbers reflect N_f_ (with ancestral N_f_ at the point of the triangle and current N_f_ at the base of the red triangle), blue numbers reflect ancestral and current N_m_, the numbers in the black oval indicate the founding effective sizes for the intial out-of-Africa migration, and dates on arrows indicate divergence times based on the tree in **Supplementary Figure 14**. Arrows are meant to indicate the schematic direction of migrations and should not be taken as indicating literal migration pathways, e.g. the results indicate divergence of the ancestors of Oceanians 61,000 years ago, but not the route(s) people took to get to Oceania.

Previous studies of N_f_ and N_m_ have largely relied on comparisons of X chromosome vs. autosomal variation, and have come to contradictory conclusions concerning the historical N_f_/N_m_ ratio^8, 9, 30^, because of methodological differences, difficulties in accounting for differences in male vs. female mutation rates, as well as the potentially greater effect of selection on the X chromosome^10, 11^. Comparison of mtDNA vs. NRY variation offers a more direct assessment free of some of the issues concerning X:autosome comparisons, but requires unbiased estimates of NRY variation, which until our study were only available from either whole genome sequencing studies^5, 12–14^ or more limited targeted studies of NRY sequence variation^7, 31^. Our results support a consistent excess of N_f_ vs. N_m_ starting even before the out-of-Africa migration that has been carried through almost all subsequent migrations (with the possible exception of East Asia), and has become even more pronounced in recent times (**Supplementary Figure 16**; **Figure 4**) due to higher rates of growth in N_f_ than in N_m_ (**Figures 3** and **4**).

In conclusion, we have developed a rapid and cost-effective means of obtaining unbiased NRY sequence information at comparable resolution to that of complete mtDNA genome sequences. Application to the HGDP provides new insights into the comparative demographic history of males and females, including support for larger between-population differences for the NRY than for mtDNA (albeit with considerable regional variation), significant bottlenecks associated with the migration of modern humans out-of-Africa and with other migration events; and overall higher female than male effective population sizes. We anticipate that this approach should enable more detailed comparative analyses of the demographic history of males vs. females, and the influence of sex-specific processes during human evolution.

## METHODS

Methods and any associated references are available in the online version of the paper.

*Note: Supplementary information is available in the online version of the paper*.

## ACKNOWLEDGMENTS

We thank: Maanasa Raghavan and Martina Codina for technical assistance; the MPI-EVA Evolutionary Genetics Sequencing Group and Bioinformatics Group for carrying out the sequencing and the initial processing of the sequence data; and Chiara Barbieri, Peter de Knijff, Brigitte Pakendorf, Kay Prüfer, and Udo Stenzel for helpful discussion. Research supported by the Max Planck Society; we acknowledge financial support from the Biotechnology and Biological Sciences Research Council (to Francois Balloux).

## AUTHOR CONTRIBUTIONS

M.S. designed the study. S.L. designed the NRY capture microarray. A.B., S.L. and R.S. carried out the laboratory work. S.L., M.L. and G.R. processed the sequences.

S.L., H.X., and A.K. analyzed the data. M.S. and S.L. wrote the paper, with input from all authors.

